# Microbial community dynamics of surface water in British Columbia, Canada

**DOI:** 10.1101/719146

**Authors:** Miguel I. Uyaguari-Diaz, Matthew A. Croxen, Kirby Cronin, Zhiyao Luo, Judith Isaac-Renton, Natalie A. Prystajecky, Patrick Tang

**Author notes:** Department of Microbiology, University of Manitoba, Winnipeg, Manitoba, Canada R3T 2N2 (Miguel I. Uyaguari-Diaz). Provincial Laboratory for Public Health, Edmonton, Alberta, Canada T6G 2J2 (Matthew A. Croxen). Department of Laboratory Medicine and Pathology, University of Alberta, Edmonton, Alberta, Canada, T6G 2R7 (Matthew A. Croxen). Public Health Ontario Laboratories, Toronto, Ontario, Canada M5G 1M1 (Kirby Cronin). National Microbiology Laboratory, Public Health Agency of Canada, Winnipeg, Manitoba, Canada R3E 3R2 (Kirby Cronin). British Columbia Cancer Agency, <u>Provincial Health Services Authority</u> Vancouver, Canada V5Z 4E6 (Zhiyao Luo). Department of Pathology, Sidra Medicine, Doha, Qatar (Patrick Tang). Department of Pathology and Laboratory Medicine, Weill Cornell Medicine, Doha, Qatar (Patrick Tang). These authors contributed equally to this work (Miguel I. Uyaguari-Diaz, Matthew A. Croxen).

## Abstract

Traditional methods for monitoring the microbiological quality of water focus on the detection of fecal indicator bacteria such as *Escherichia coli*, often tested as a weekly grab sample. To understand the stability of *E.coli* concentrations over time, we evaluated three approaches to measuring *E. coli* levels in water: microbial culture using Colilert, quantitative PCR for *uidA* and next-generation sequencing of the 16S rRNA gene. Two watersheds, one impacted by agricultural and the other by urban activities, were repeatedly sampled over a simultaneous ten-hour period during each of the four seasons. Based on 16S rRNA gene deep sequencing, each watershed showed different microbial community profiles. The bacterial microbiomes varied with season, but less so within each 10-hour sampling period. *Enterobacteriaceae* comprised only a small fraction (<1%) of the total community. The qPCR assay detected significantly higher quantities of *E. coli* compared to the Colilert assay and there was also variability in the Colilert measurements compared to Health Canada’s recommendations for recreational water quality. From the 16S data, other bacteria such as *Prevotella* and *Bacteroides* showed promise as alternative indicators of fecal contamination. A better understanding of temporal changes in watershed microbiomes will be important in assessing the utility of current biomarkers of fecal contamination, determining the best timing for sample collection, as well as searching for additional microbial indicators of the health of a watershed.

## Introduction

The use of *Escherichia coli* as a fecal bacterial indicator in water is widely-adopted but also contentious (Ferguson & Signoretto 2011; Isobe et al. 2004; Rochelle-Newall et al. 2015). As a species, *E. coli* is extremely diverse in its ability to colonize a wide range of hosts and in the range of diseases it can cause (Croxen & Finlay 2010; Kaper et al. 2004). Since *E. coli* colonizes the gastrointestinal tract of most birds and animals, and is readily culturable from feces, it seems to be well suited as an indicator of fecal contamination. Many regulatory bodies around the world use *E. coli* as an indicator of water quality and the amount of *E. coli* cultured from water is used as an actionable metric to enact decisions on boil water advisories or beach closures. A major problem with *E. coli* as an indicator organism, however, is the limited amount of information that can be derived from a positive- or negative-culture result. While a positive result may be indicative of recent fecal pollution or also represent environmental re-growth from non-recent fecal deposition, it does not provide information on the source of contamination or on the risk to health. A negative result does not rule out other pathogens such as viruses or protozoa, exemplified by an outbreak in Wisconsin, where over an estimated 400,000 people fell ill due to *Cryptosporidum parvum* oocysts present in municipal waters; this despite treated water samples testing negative for coliforms (Mac Kenzie et al. 1994). Furthermore, while the ability of *E. coli* to survive in the environment appears poor under simulated conditions (van Elsas et al. 2011), there have reports of *Escherichia* spp. naturally present in the environment (phenotypically identical to *E. coli*) (Byappanahalli & Fujioka 1998; Clermont et al. 2011; Walk et al. 2009). These issues raise questions about the usefulness of *E. coli* for routine water quality testing purposes.

Routine water quality testing involves the collection of a water sample (“grab sample”) and culturing it for a fecal bacterial indicator such as *E. coli*. Culture-positive results are then reported either as a single maximum concentration or as a geometric mean over a defined sampling period (BC 2017). How representative one result, from one sampling event or averaged from previous week’s results, is uncertain. Culture of two separate 100 ml water samples collected at the same time may yield different *E. coli* results (Wohlsen et al. 2006). Whether this is merely representative of a minor, culturable community member, such as *E. coli*, or true for the entire community, is not yet known.

It is important to protect watersheds as deterioration of their health impacts the environment and inter-connected ecosystems. A large metagenomics project was carried out to develop biomarkers to better measure water health compared to current methods (www.watersheddiscovery.ca). The work presented here provides a foundation to better understand the dynamics of source water microbial communities over the course of a day, and during four quarters throughout the year.

## Materials and methods

### Water sample collection

Two source waters were chosen from the lower mainland area of British Columbia. The first site is a slough that runs through dairy, poultry and produce farms and is impacted by agricultural land use. The second site is impacted by urban land use; it runs through an urban development and includes parkland that is used for recreational purposes. Sampling days were on January 29^th^, 2013 (Quarter 1; Q1), April 29^th^, 2013 (Q2), July 31^st^, 2013 (Q3), and October 29^th^, 2013 (Q4) with typical weather patterns for the area. The first set of samples was collected at 08:00 Pacific Standard time (PST), followed by a second set collected 10 minutes later (08:10 PST). Collection of water in this manner occurred every two hours until 18:10 PST. At each time point, four samples were collected in succession; two 250 ml samples were collected for filtration in 250 ml wide mouth translucent HDPE bottles. Two additional samples were collected for Colilert (IDEXX, Westbrook, ME) in narrow mouth 250 ml high-density polyethylene bottles containing sodium thiosulfate (Systems Plus, Baden, ON). Samples were kept on ice throughout the day, stored at 4 °C overnight, and processed within 24 hours. In total, 48 samples (24 for filtration and 24 for Colilert) were collected at each site, per day for an overall total of 196 filtration samples (24 samples × 2 sites × 4 sampling days) and 196 Colilert samples.

### Water filtration, coliform and *E. coli* counts

One hundred milliliters (ml) of water were filtered using 0.2-μm 47-mm Supor-200 polyethersulfone membrane disc filters (Pall Corporation, Ann Harbor, MI) inserted into 300 ml Pall 47 mm magnetic polyphenylsulfone filter funnels (Pall Corporation, Ann Harbor, MI). The filters were rolled up and stored at -20°C until needed for nucleic acid extraction. A negative filtration control was also processed using MilliQ water (Millipore Corporation, Billerica, MA). Dilutions were conducted from the bottles containing sodium thiosulfate (1:10 for the urban site, 1:100 for the agricultural site) and 100 ml of diluted sample was poured into a Colilert Quanti-Tray/2000 (IDEXX). Trays were sealed according to manufacturer’s instructions and incubated at 37°C. Yellow (coliforms) and blue fluorescent wells (*E. coli*) were enumerated after 24 hours of incubation, and used to calculate the most probable number (MPN) according to the table provided with the Colilert kit. MilliQ water was used as a negative control.

### Nucleic acid extraction, 16S rRNA gene amplification and sequencing

Following filtration polyethersulfone membrane disc filters were cut into four pieces. Nucleic acids were extracted from each quarter filter using the MO BIO PowerLyzer PowerSoil DNA Extraction Kit (MoBio, Carlsbad, CA) according the manufacturer’s instructions. Extracted DNA samples from each quarter filter were pooled and further concentrated using 0.1 volumes of 3 M sodium acetate, two volumes of 100 % ethanol, and 5 μl of 5 μg/μl linear acrylamide. Eluents were stored at –80 °C overnight, and then centrifuged at 17,000×g for 30 min at 4 °C. Supernatants were discarded, and pellets were washed with 70 % ice-cold ethanol, air dried, and resuspended in 34 μl of 10 mM Tris, pH 8.5. One µl of DNA from each extract was used to resuspended in 34 μ amplify the V4 region (515F: 5’-GTGCCAGCMGCCGCGGTAA-3’ and 806R: 5’-GGACTACHVGGGTWTCTAAT-3’) of the 16S ribosomal RNA (ca. 253 bp) using GoLay primers as previously described (Caporaso et al. 2012). These GoLay primers add compatible adapters and barcodes for the Illumina MiSeq and HiSeq sequencing platforms. The negative processing control was also included in the PCR amplification, as well as a mock community that consisted of *E. coli, Pseudomonas putida, P. aeruginosa, P. fluorescens, Burkholderia cenocepacia, Bacillus amyloliquefaciens, B. cereus, Rhodobacter capsulatus, Streptomyces coelicolor, Micrococcus luteus*, and *Frankia sp*. CcI3 as described earlier (Peabody et al. 2015).

Amplicons were purified using the QIAgen QIAquick PCR purification kit according to manufacturer’s instructions. Purified PCR products were quantified using Qubit dsDNA HS Assay Kit (Invitrogen). Each amplicon from a single sampling day (24 agricultural, 24 urban, 1 negative control and, 1 mock community) was pooled together for a final equimolar concentration of 4 nM. Each pool (4 total) was diluted to a final loading concentration of 11.5 pM, and PhiX at 8 pM was spiked in at 30%. Each pool was run individually on an Illumina MiSeq (Illumina, Inc., San Diego, CA) using the 300-cycle (2 × 150 bp) MiSeq Reagent Kit v2 (Illumina). Immediately prior to sequencing, additional primers (100 µM) were added to the MiSeq reagent cartridge as follows: 3.4 µl of Read 1 sequencing primer to reservoir 12; 3.4 µl of Index sequencing primer to reservoir 13; 3.4 µl of Read 2 sequencing primer to reservoir 14 (Caporaso et al. 2012).

### Quantitative PCR for *E. coli*

Quantitative PCR (qPCR) was used to target the β-glucuronidase gene (*uidA*) of *E. coli* and *Shigella* spp., and amplified an 84 bp product. The oligonucleotides 784F (5’-GTGTGATATCTACCCGCTTCGC-3’), 866R (5’-GAGAACGGTTTGTGGTTAATCAGGA-3’) and TaqMan probe EC807 (5’-FAM-TCGGCATCCGGTCAGTGGCAGT-BHQ1-3’) have been previously described (Frahm & Obst 2003), except TAMRA was replaced with the BHQ1 quencher. All oligonucleotides were purchased from Integrated DNA Technologies (IDT, Inc., Coralville, IA).

Each qPCR reaction was done in a 20 µl volume which consisted of 2 µl of template, 1X PerfeCTa qPCR ToughMix, UNG, Low ROX (Quanta), 0.4 µM 784F, and 0.4 µM 866R. Template DNA was diluted 10-fold for agricultural samples, while template DNA from urban samples were not diluted. Amplification was performed on MicroAmp Fast Optical 96-Well Reaction Plates (Life Technologies, Carlsbad, CA) on an Applied Biosystems 7500 Real-Time PCR System (Applied Biosystems). The following cycling conditions were used: 45 °C for 5 minutes, 95 °C for 3 minutes, then 40 cycles of 95 °C for 15 seconds and 60 °C for 1 minute. A standard curve was included in each qPCR run on 10-fold serial dilutions (1,740,000 to 174 copies) of genomic DNA from *E. coli* ATCC 25922 that was extracted using the QIAgen QIAamp DNA Mini Kit (Qiagen Sciences, Inc., Germantown, MD) according to the manufacturer’s protocol. Each sample was run in duplicate, while standards were run in triplicate. Gene copy numbers were normalized per 100 ml of sample.

### 16S rRNA gene qPCR assay

To estimate bacterial copy number in water samples, copy numbers of a 16S rRNA gene fragment (∼352 bp) of were calculated as described by Ritalahti et al. (2006). Primers Bac1055YF (5’-ATGGYTGTCGTCAGCT-3’) (Ferris et al. 1996; Ritalahti et al. 2006) and Bac1392R (5’-ACGGGCGGTGTGTAC-3’) (Lane 1991) were used in combination with Probe Bac1115 containing a 5’ 6-FAM dye (CAACGAGCGCAACCC) (Harms et al. 2003; Lane 1991) with an internal ZEN quencher and a 3’ Iowa Black fluorescent quencher (Life Technologies, Carlsbad, CA). Due to multiple copy nature of the 16S rRNA gene in a bacterium, copy numbers were normalized by a factor of 4.3 (Lee et al. 2009) per 100 ml of sample. *E. coli* genomic DNA l real-time PCR reaction consisted of was used for standard curves for *16S rRNA* gene. Each 20 μl real-time PCR reaction consisted of 10 μl of ABI TaqMan universal master mix (Life Technologies, Carlsbad, CA), 0.4 μM of each primer, 0.1 μM of probe and template DNA. DNA samples from agricultural site were diluted primer, 0.1 μ l of 1:10 using nuclease free-water (Promega Corporation, Fitchburg, WI), while that 1 to 2 μl of undiluted template DNA were used for urban water samples, and 1 μl. Quantitative PCR reactions were conducted on an Applied Biosystems 7500 Fast real-Time PCR system (Life Technologies, Carlsbad, CA). The thermal cycling conditions consisted of incubation for 2 min at 50 °C, initial denaturation for 10 min at 95 °C followed by 40 cycles of 15 s at 95 °C and 60 s at 60 °C. Standards were run in triplicate, while that environmental samples were run in duplicate.

### Sequence analysis

Demultiplexed forward and reverse reads were error corrected using BayesHammer (Nikolenko et al. 2013), followed by adapter and primer trimming, and quality filtering using Trimmomatic v0.32 (Bolger et al. 2014). Trimmed and filtered reads were assembled using PANDAseq v2.2 (Masella et al. 2012), and any assembled sequences under 220 nucleotides were discarded. The QIIME v1.9.1 package (Caporaso et al. 2010b) was used to compare the microbial communities based on 16S sequences. We used UCLUST (Edgar 2010) to pick Operational Taxonomic Units (OTU) by following a workflow for open-reference clustering (Rideout et al. 2014) at 97% using the GreenGenes v13_8 database (DeSantis et al. 2006). After OTU picking, OTUs with abundances less than 0.005% were removed as recommended by Bokulish et al. (Bokulich et al. 2013). ChimeraSlayer (Haas et al. 2011) was used to identify chimeras and the remaining OTUs were aligned using PyNast (Caporaso et al. 2010a) before a phylogenetic tree was constructed using FastTree2 (Price et al. 2010). Bray-Curtis dissimilarity was used to determine beta diversity for each sample. Figures were generated with QIIME, R (ggplots2) (Wickham 2009), and hclust2 (https://bitbucket.org/nsegata/hclust2).

### Data analysis

All data were log_10_ transformed for analysis. Spearman’s rank correlation analysis was conducted among variables. Paired t-tests were used to assess differences between qPCR and Colilert assays. Data was analyzed using Statistical Analysis System (SAS, version 9.1 for Windows). A p-value of 0.05 was assumed for all tests as a minimum level of significance.

### Data availability

All sequences generated for this study are available under NCBI BioProject PRJNA287840, SRA samples SRR2083863 through SRR2084062.

## Results and discussion

### Sampling sites

The lower mainland of British Columbia (BC) has temperate weather. We chose two study watersheds that are impacted differently; one is impacted by farming land use (Agricultural; AGR), and one has more anthropogenic impact through proximity to residential and recreational land use (Urban; URB). Fig 1 represents AGR and URB sites with land cover. Distance between these non-connected watersheds is ∼65 Km.

**Fig 1.**
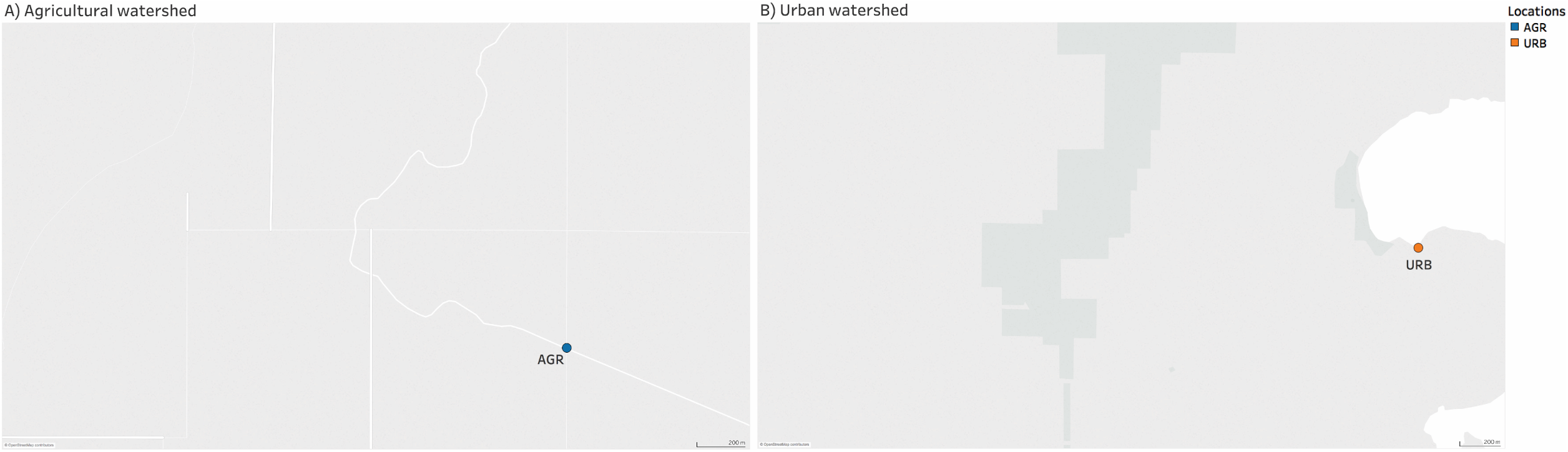
Sampling locations with land cover. A) Agricultural watershed. B) Urban watershed

### *E. coli* counts in watershed samples can range depending on sampling event and time

We collected surface water samples during the winter (Q1), spring (Q2), summer (Q3) and fall (Q4). We recognized that due to the single sampling event, these results are still a snapshot in that particular quarter of the year and should not be interpreted as representative of an entire season, or even reflective of a typical season in the study area. As well, we collected samples during a defined daily period. Water samples for assessing water quality in British Columbia are typically collected once a week for *E. coli* culture. To also assess the variability of *E. coli* counts in the agricultural- and urban-impacted watershed study sites, we collected samples in duplicate, followed in duplicate again 10 minutes later every two hours, from 08:00 hrs until 18:00 hrs. Agreement between duplicate samples varied; sometimes samples did not differ, when other times they did. The most extreme example of this were the results of Agricultural Q1 at the 16:10 hrs time point. Sample A showed 100 *E. coli* MPN/100 ml, whereas the duplicate sample (B) had 12X more culturable *E. coli* cells (Fig 2). Similarly, *E. coli* counts fluctuated during the different time points during the day in both the agricultural and urban watersheds. Health Canada’s recommendations for recreational water quality is 400 *E. coli* MPN in 100 ml from a single sampling event (Canada 2012; Levesque & Gauvin 2007). Some *E. coli* counts were variable during the day and there are many instances where the sampling time would have made a difference to the interpretation and concomitant action of public health officials based on Health Canada’s guidelines (red dotted line; Fig 2). Because of sampling location and transportation to the laboratory, we acknowledge sample holding times exceeded the preferred time interval of 8 hours from collection to examination (APHA 2005; Bartram & Rees 2000). In the present study, samples were processed within 24 hours. No significant differences in counts of *E. coli* and total coliforms have been reported in water samples up to 27 hours since collection (Aulenbach 2010; Maier et al. 2015; Pope et al. 2003).

**Fig 2.**
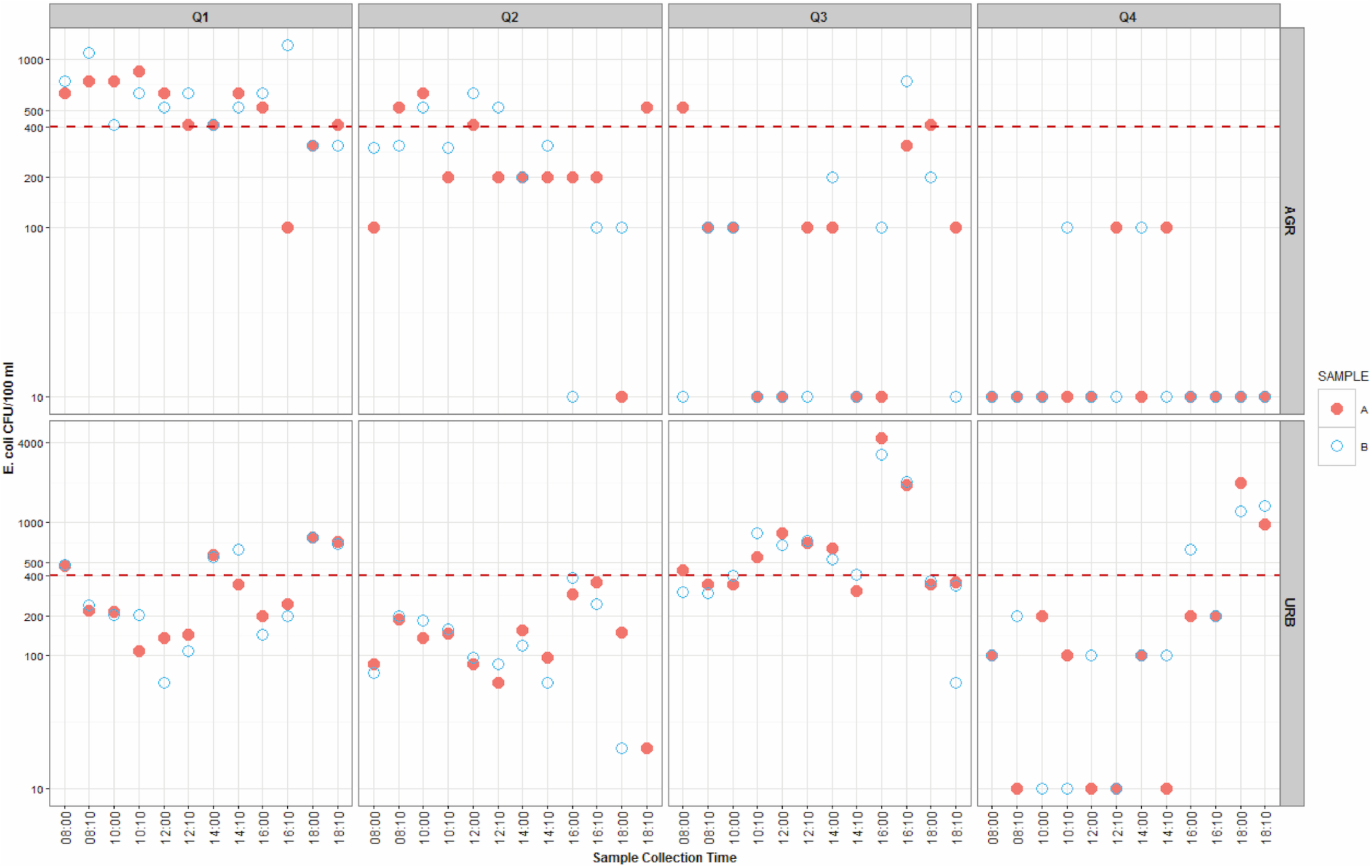
Scatterplot of *E. coli* colony counts using Colilert. The x-axis represents sample collection time in each quarter (Q1-Q4) and the y-axis depicts the *E. coli* most probable number (MPN) per 100 ml of water sample. Full red and empty blue circles depict true sample replicates. Red dotted line set at 400 MPN/100 ml represents Health Canada’s limit for recreational water quality.

### Comparison of *E.coli* counts and *uidA* gene quantitative PCR assay

We then compared the number of viable *E. coli* cells to the single copy *uidA* gene, detected by qPCR. There was a moderate positive correlation between the culturable *E. coli* and the qPCR results (*r*_*s*_= 0.5374, p-value<0.0001), suggesting that both methods shared an agreement of 28.9% of common values. This comparison also indicated a difference of approximately 2 orders of magnitude in test results, with higher numbers detected by the *uidA* gene qPCR assay (p-value<0.0001). While the *uidA* gene is specific for *E. coli* and some *Shigella* spp. (Maheux et al. 2011), this qPCR assay detects DNA from viable, viable but non-culturable, as well as dead cells. When compared to other studies (Noble et al. 2010; Oliver et al. 2016), this association seems to be low, however a drop in correlation coefficients have been reported in environmental water samples (Walker et al. 2017).

### Microbial community diversity differ in the long-term, but not short-term

Since *E. coli* counts varied throughout the day, we wanted to examine the dynamics of bacterial communities. We used the same water samples to conduct deep amplicon sequencing. In agricultural sites and across seasons, clusters clearly stood out from each other suggesting a more or less steady microbial community all day long (Fig 3A). While clusters were also found in urban sites, these data points were more spread out within the groups with some overlap in the community composition (Fig 3B). Factors such as water use, runoff, possible leaking of wastewater or seepage (Vermonden 2010; Vermonden et al. 2009) may explain such perturbations (Fig 3B). Results from the 16S rRNA gene sequencing and top 50 OTUs at the family level are depicted in Fig 4. *Enterobacteriaceae* constituted a relatively small fraction (<1%) of the total microbial community. Relative abundance values of 0.06% and 0.76% were observed for this family in AGR (Fig 4A) and URB (Fig 4B), respectively. Although *E. coli* was detected using both the *uidA* qPCR and testing methods, it could not be detected using deep amplicon sequencing. This observation, not only for *E.coli* but also for other sequences, may be related to amplicon fragment size (254 bp). This may be a limitation to obtaining sufficient genetic information (Calus et al. 2018) to identify bacteria at the genus or species level.

**Fig 3.**
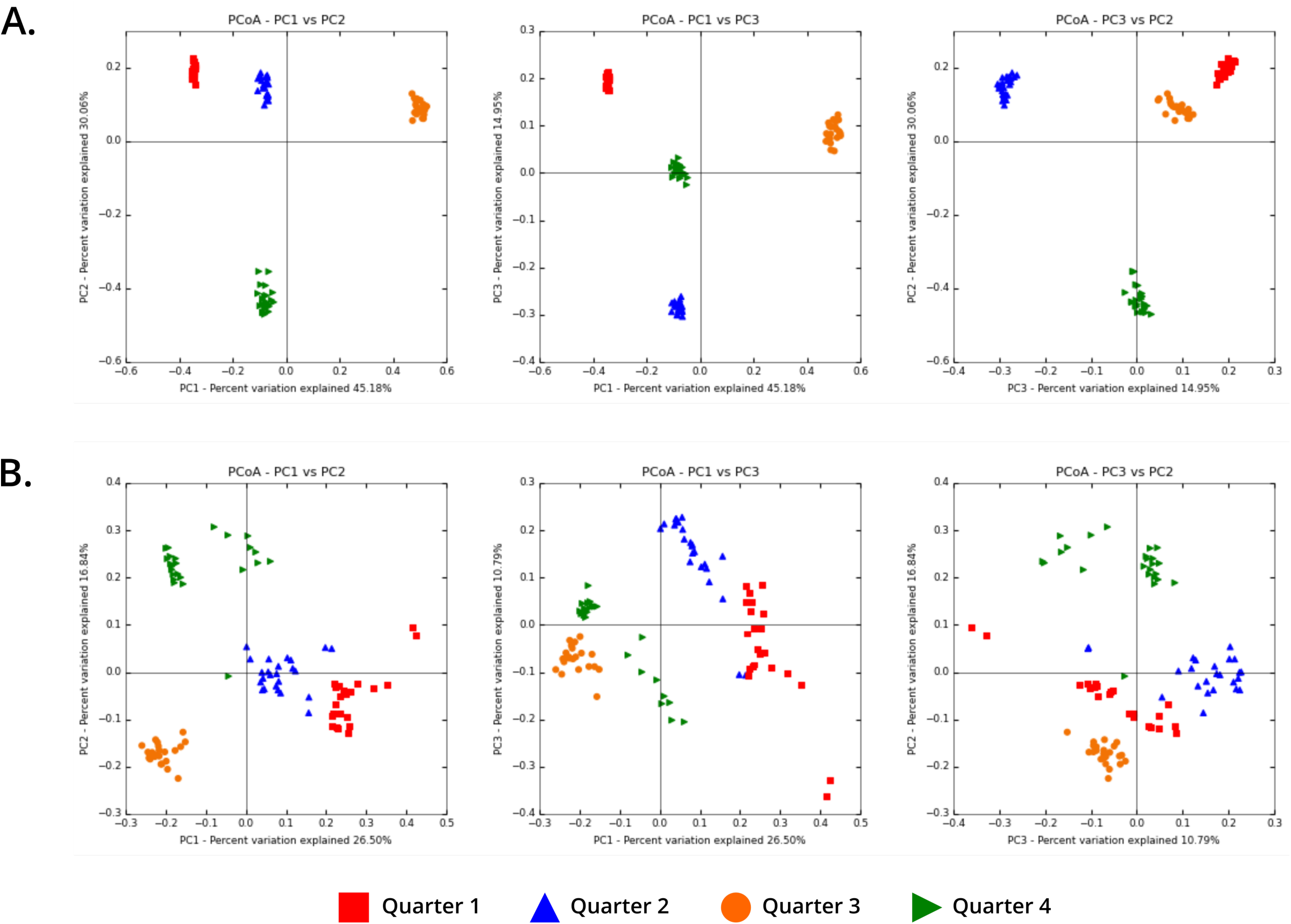
Principal coordinate analysis based on Bray-Curtis dissimilarity (β-diversity) for the: β 16Sr RNA gene amplicons (n=24 time points) across seasons (Q1-Q4). A) Agricultural watershed. B) Urban watershed.

**Fig 4.**
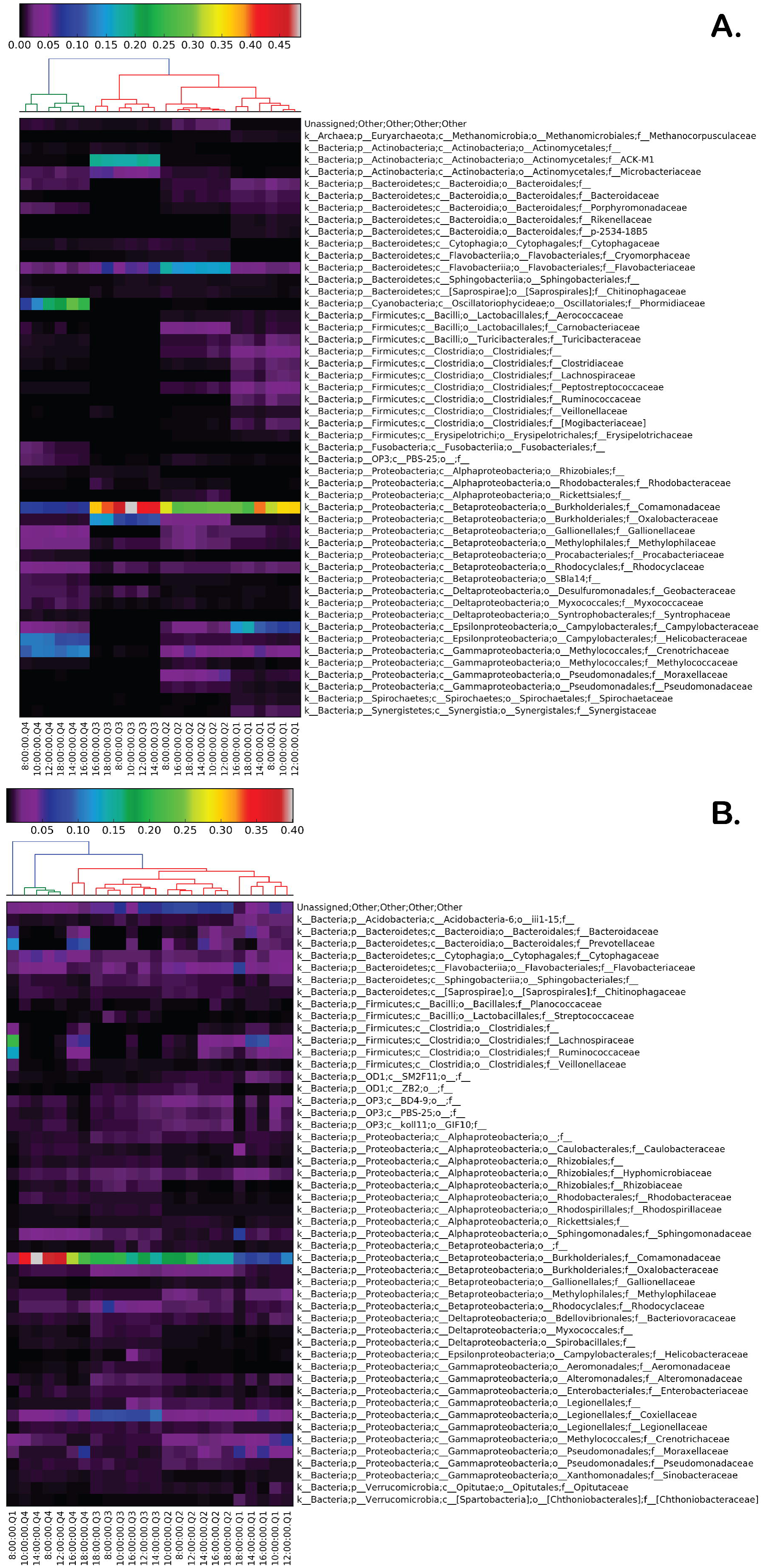
Heat map depicting the relative abundance of the top 50 most abundant OTUs in agricultural (A) and urban (B) watersheds. Hourly data sets were merged using QIIME, based on categorical data for visualization purposes. The color key at the top represents percentages of relative abundance, while that branches depict hierarchical clustering based on similarity.

In the agricultural study site, two clusters were observed for bacterial communities between Q4 (fall) and Q3 (summer), Q2 (spring) and Q1 (winter). Within this latter group, 3 sub-groups were identified. In both sampling sites, the most abundant OTUs were in the family *Comamonadaceae* with average values across seasons of 28.1% and 18.9% for AGR and URB, respectively. Similar estimates have been reported for this family in aquatic systems impacted by urbanization or agricultural activities (Griffin et al. 2017; Lopes et al. 2016; Newton & McLellan 2015; Willems 2014). Although *Comamonadaceae* was predominant across seasons, different patterns were observed for this family. For example, in AGR, a higher abundance of *Comamonadaceae* was observed during Q3, while members of this family were more abundant during Q4 in URB. A possible explanation for these patterns is the manure application occurring on study site farms during this part of the year (summer) and weather related runoff into the sampling site. In URB, a higher abundance of *Comamonadaceae* was observed in Q4, corresponding to the onset of the rainy season with storm water runoff in the region. While not considered human pathogens, two genera (*Xylophilus* and some species of *Acidovorax*) within this family affect plants (Willems 2014). Other relevant major families (≥4%) observed in AGR were *Flavobacteriaceae* (6.7%), *Crenotrichaceae* (5.4%) and *Campylobacteraceae* (4.3%). Members of the first two families are widely distributed in the environment (McBride 2014; Siljanen et al. 2011; Urios et al. 2006), with similar estimates being reported in agricultural settings (Pandey et al. 2018; Shawkey et al. 2009) and water-sediment interfaces (Chidamba & Korsten 2015; Frindte et al. 2016). On the other hand, some members of *Campylobacteraceae* such as *Campylobacter* and *Arcobacter* contain species that are well-known human pathogens (Lastovica et al. 2014; Lehner et al. 2005). Although *Campylobacteraceae* was detected in both sites, we observed an average of ten-fold difference between AGR (4.1%)(Fig 4A) and URB (0.4%) (data not shown). Interestingly, the relative abundance of *Campylobacteraceae* in AGR was at least twice as high during Q1 (winter) compared to the other seasons (Fig 4A), with no major changes observed in URB. Note that members of *Campylobacteraceae* are livestock and poultry associated (Mughini Gras et al. 2012), both of which are farmed in the study area. This *Campylobacteraceae* prevalence, concentration and survival during winter compared to other seasons in AGR, may be associated to with their sensitivity to desiccation during drier months (Smith et al. 2016), occurrence of heavy rainfall (Ahmed et al. 2013; Moriarty et al. 2011), their presence in waterfowl feces (Moriarty et al. 2012), or various farming practices (Rapp et al. 2014). Alternatively this could also represent detection of material from bacterial lysis (Feng et al. 2017). Another family observed with high relative abundance (10-30%) in AGR was *Phormidiaceae* but this family was only abundant during fall (Q4), and completely absent or in very low abundance during other seasons (Fig 4A). Members of this family are common in poor quality streams with precipitation triggering its bloom during fall (Hossain et al. 2012; Kaestli et al. 2017).

It is important to mention that in URB fecally-associated bacteria such as *Prevotella* (2.20%), *Bacteroides* (1.29%), *Blautia* (0.76%), *Faecalibacterium* (0.57%) and *Coprococcus* (0.24%) were observed (Figs 4B and 5). These genera have been proposed as potential indicators of sewage and human fecal contamination (Eren et al. 2015; Fisher et al. 2015; Fogarty & Voytek 2005; McLellan et al. 2013). Significant positive correlations were detected among these genera (*r*<0.5137, p<0.001). Major peaks on these genera of bacteria in URB occurred in Q1 during 8 AM and 2 PM (PST) (Fig 5). Moreover, a relatively high abundance of these genera was observed during Q2, Q3 and Q4 between 2 PM and 4PM. Besides precipitation (17.8 mm) in URB during Q1, these observations may represent water use (i.e. flushing events) occurring during the day; with the latter time reflecting changes associated to daylight saving time. When compared to the agricultural watershed, *Prevotella, Bacteroides*, and *Coprococcus* were present in AGR, but in combination these genera made up less than 0.23% of the total microbial community. We observed a positive correlation between *Prevotella* and *Bacteroides* (*r<* 0.7479, p<0.001). Genera such as *Blautia* and *Faecalibacterium* were not detected in AGR using with deep amplicon sequencing.

**Fig 5.**
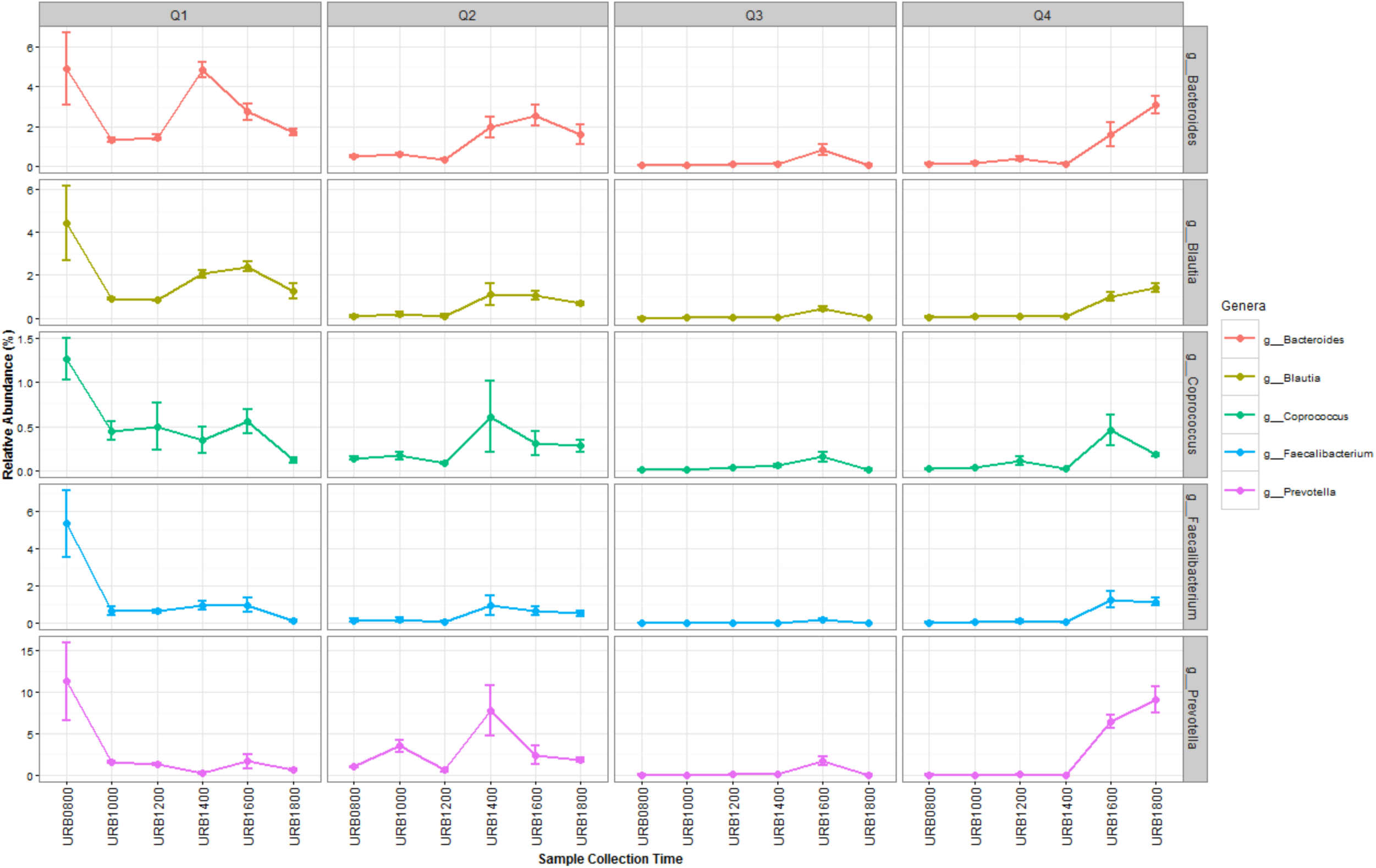
Line plots of fecally-associated bacteria observed in the urban watershed. The x-axis represents sample collection time in each quarter (Q1-Q4) and the y-axis shows percentages of relative abundance.

The prevalence of *Prevotella* and *Bacteroides* in URB may also reflect biomarkers of the sampled urban setting (Gorvitovskaia et al. 2016). Another relevant taxa reported to be part of the human gut microbiome (Arumugam et al. 2011) and observed in our study was *Ruminococcus*. However, low average values of 0.07% and 0.10% were found for AGR and URB, respectively. The fact that no major differences between sites were observed across seasons for *Ruminococcus* may indicate either an anthropogenic influence occurring in AGR. We also cannot rule out wildlife as contributor of this genera (Arumugam et al. 2011). Overall, genera such *Prevotella* and *Bacteroides* were present in higher relative abundance in URB compared to the other fecally associated bacteria. These two genera have been proposed as alternative indicator to assess water quality impacted by anthropogenic activities (Fogarty & Voytek 2005).

To potentially identify novel biomarkers of water quality more detectable than *E.coli*, additional correlation analyses were conducted with the 10 more abundant bacterial families in each watershed and physico-chemical water quality parameters (Fig 6). In AGR and URB sites there was a moderate to strong positive correlation among variables such as 16S rRNA, *uidA* gene, *E. coli* colonies and total coliforms (as measured by Colilert) (Fig 6). Among these variables, 16S rRNA gene copy numbers were positively correlated with *Comamonadaceae*, the most abundant bacteria detected in AGR and URB by deep amplicon sequencing. Moreover, water quality parameters related with conductivity (specific conductivity, salinity, total dissolved solids, and oxidation-reduction potential) were positively (p<0.0001) correlated with *Comamonadaceae*. In contrast, this family was negatively correlated with dissolved oxygen (DO). This distinguishable pattern has also been reported for *Comamonadaceae* in anthropogenic impacted aquatic systems (Aguirre et al. 2017).

**Figure 6.**
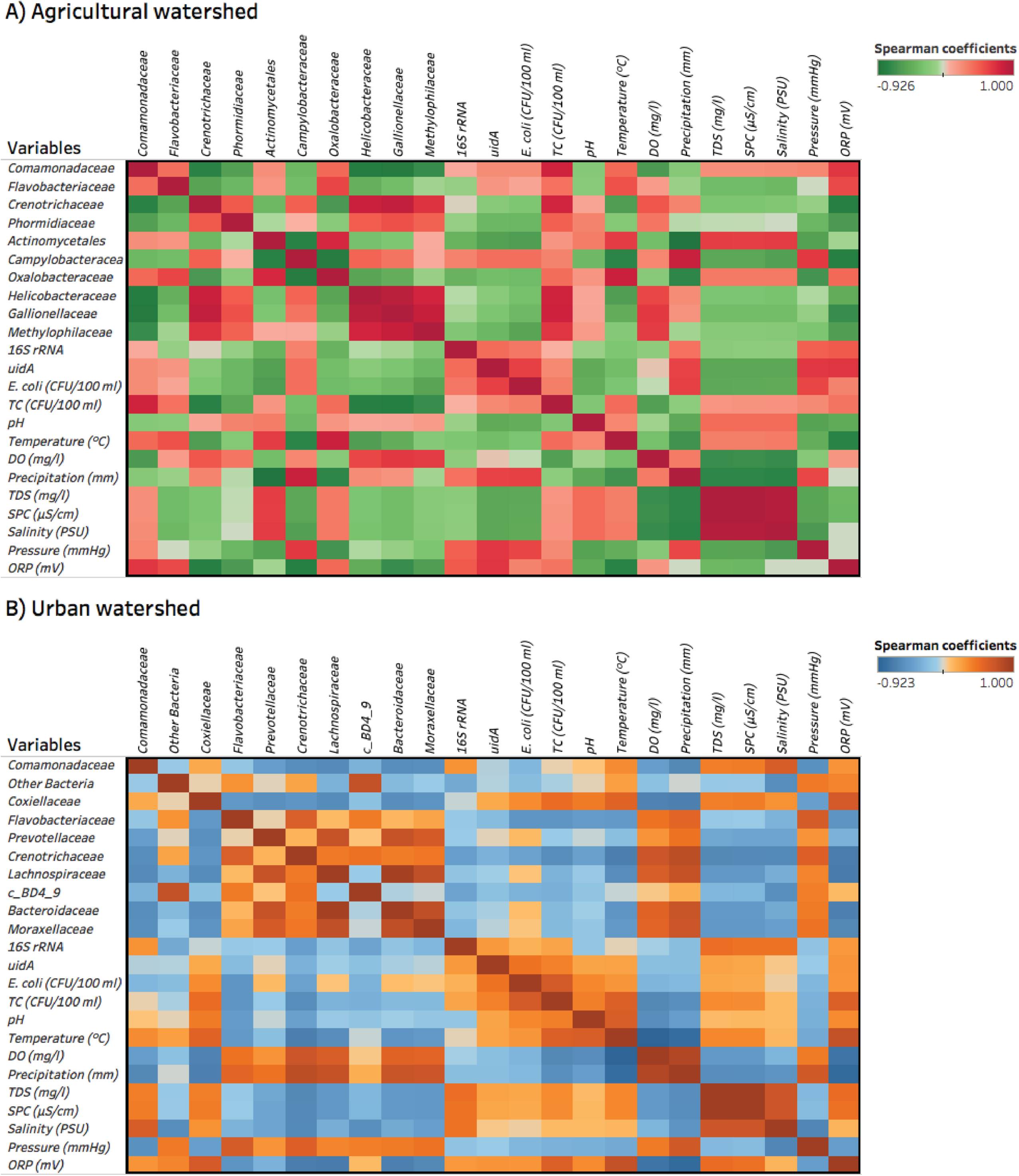
Heat map showing the Spearman’s rank correlation analysis between the top 10 most abundant families and water quality parameters. A) Agricultural watershed. B) Urban watershed.

Temperature had a positive correlation with *Comamonadaceae* and a negative correlation with *Crenotrichaceae* in both watersheds. In AGR, pH was significantly correlated with seven of the most abundant families including *Crenotrichaceae, Phormidiaceae*, Actinomycetales, *Oxalobacteraceae, Helicobacteraceae, Gallionellaceae, Methylophilaceae*. Moderate changes in pH may cause community shifts especially in less abundant groups of bacteria or pH-susceptible groups (Krause et al. 2012) as the ones described in our study. Similar to the observations reported by Krause et al. 2012, pH did not have a significant effect on the overall bacterial abundance in both watersheds as estimated by the 16S rRNA gene (Fig 6).

Dissolved oxygen and precipitation were parameters highly correlated between them in both watersheds. This association was reflected by members of the *Crenotrichaceae* family in AGR and URB. Other families associated to these 2 variables and mostly positively associated between them included *Flavobacteriaceae, Prevotellaceae, Lachnospiraceae*, a proposed novel bacterial division BD4-9 (possibly from the phylum Verrucomicrobia) (Briee et al. 2007; Derakshani et al. 2001), *Bacteroidaceae* and *Moraxellaceae* in URB. These latter two families were positively and strongly correlated between each other in URB (r_S_=0.8103, p-value<0.0001) and AGR (r_S_=0.8319, p-value<0.0001). Additional positive associations between bacteria from fecal origin in URB such as *Lachnospiraceae, Bacteroidaceae* and *Prevotellaceae* were also determined by correlation analysis (Fig 6B), and may reflect the microbiome of human populations and in particular, sewage (McLellan et al. 2013; Newton & McLellan 2015). Although multiple correlations were observed in the present study, we only described relevant associations that governed the most abundant microbes in AGR and URB (Fig 6).

## Conclusions

Metagenomic analysis of bacterial communities indicated fairly stable patterns, while *E. coli* counts varied widely over time. A moderate correlation was detected between culturable *E. coli* (as measured by Colilert) and *uidA* gene. Nevertheless, a difference of 2 orders of magnitude was observed between these approaches. The structure of microbial communities did not change significantly on an hourly basis, however dramatic changes were observed across seasons in the two study locations. Although each watershed exhibited different microbial profile patterns, *Comamonadaceae* was the most abundant bacteria detected in both watersheds by 16S rRNA gene deep sequencing. Relative abundance of *Campylobacteraceae* indicated a ten-fold difference between AGR (4.1%) and URB (0.4%). Moreover, members of the family *Enterobacteriaceae* made up less than 1% of the total microbial community in AGR and URB. In this context, *E. coli* could not even be detected at the genus level. Genera such *Prevotella* and *Bacteroides* were present in higher relative abundance in URB compared to other fecally associated bacteria, which may suggest alternatives in the search for better biomarkers of fecal pollution. Precipitation and DO were positively correlated with seven of the most abundant bacterial families in URB, while that in AGR, pH was positively correlated with the same number of families. Besides a taxa-based approach such as 16S rRNA gene sequencing, the use of metagenomics may be an opportunity to discover better fecal pollution biomarkers. Discovery approaches could include use of taxonomic information but also information at the metabolic and functional level (Simon & Daniel 2011). Although it has been demonstrated that microbial communities may change over short distances (10 km) (Hewson et al. 2006), we found that the study watersheds showed generally stable community structure over a 10-hour period, but we did see notable differences comparing across seasons. As the accepted standard for measuring watershed health, it would be ideal to have a more somewhat less variable (counts differed greatly between samples collected at 10-minute intervals) signal. This supports our belief that biomarkers other than *E. coli* should be sought as better indicators of fecal contamination. Future studies could examine more intermediate time intervals such as sampling per week or per month, to assess possible significant changes in microbial communities at these intervals. All this information will assist in developing better methods of monitoring of water quality.

## Author Contributions

**Conceptualization:** Miguel I. Uyaguari-Diaz, Matthew A. Croxen, Natalie A. Prystajecky, Patrick Tang.

**Data curation:** Matthew A. Croxen, Miguel I. Uyaguari-Diaz, Kirby Cronin, Zhiyao Luo.

**Formal analysis:** Miguel I. Uyaguari-Diaz, Matthew A. Croxen.

**Funding acquisition:** Judith Isaac-Renton, Patrick Tang.

**Investigation:** Miguel I. Uyaguari-Diaz, Matthew A. Croxen.

**Methodology:** Miguel I. Uyaguari-Diaz, Matthew A. Croxen, Kirby Cronin, Zhiyao Luo.

**Project administration:** Judith Isaac-Renton, Natalie A. Prystajecky, Patrick Tang.

**Resources:** Natalie A. Prystajecky, Patrick Tang.

**Software:** Miguel I. Uyaguari-Diaz, Matthew A. Croxen.

**Supervision:** Judith Isaac-Renton, Natalie A. Prystajecky, Patrick Tang.

**Validation:** Miguel I. Uyaguari-Diaz, Matthew A. Croxen.

**Visualization:** Miguel I. Uyaguari-Diaz, Matthew A. Croxen.

**Writing – original draft:** Miguel I. Uyaguari-Diaz, Matthew A. Croxen.

**Writing – review & editing:** Miguel I. Uyaguari-Diaz, Matthew A. Croxen, Kirby Cronin, Zhiyao Luo, Judith Isaac-Renton, Natalie A. Prystajecky, Patrick Tang.

### Acknowledgements

We would like to thank Bridget Lee, Michael Peabody, Thea van Rossum, Alvin Xian, Mitchell Burton, Sara Tan and Tyler Nelson for help with the water sampling. We would also like to thank Dr. Fiona Brinkman (Simon Fraser University) for providing the mock community.

